# PIK3CA-related overgrowth spectrum (PROS) zebrafish models reveal pan-lineage developmental dysregulation

**DOI:** 10.64898/2025.12.03.691507

**Authors:** Hannah Brunsdon, Nuoya Wang, Micha Sam Brickman Raredon, Ralitsa R Madsen, Robert K Semple, E Elizabeth Patton

**Affiliations:** MRC Human Genetics Unit, Institute of Genetics and Cancer, The University of Edinburgh, EH4 2XU, United Kingdom; Edinburgh Cancer Research, CRUK Scotland Centre, Institute of Genetics and Cancer, The University of Edinburgh, EH4 2XR, United Kingdom; Centre for Cardiovascular Science, The University of Edinburgh, Edinburgh, EH16 4TZ, United Kingdom; Department of Anesthesiology, Vascular Biology and Therapeutics, Program in Translational Biomedicine, Yale School of Medicine, USA; MRC Protein Phosphorylation and Ubiquitylation Unit, School of Life Sciences, University of Dundee, Dundee, DD1 5EH, United Kingdom

**Author notes:** **Classification:** Biological Sciences, Developmental Biology.

**Keywords:** PIK3CA, overgrowth, vascular malformations, PI3K, zebrafish, PROS, CLOVES

## Abstract

Post-zygotic gain-of-function *PIK3CA* mutations arising during embryonic development cause disorders collectively known as the *PIK3CA*-related overgrowth spectrum (PROS). This ranges from minor, localized overgrowth to devastating multi-tissue overgrowth. Disease severity is widely attributed to a combination of PIK3CA genotype, affected cell type, and developmental timing of mutation acquisition. However, in PROS this explains neither the biased pattern of overgrowth - disproportionately affecting mesoderm and neuroectoderm-derived tissues - nor the typical low mutation burden within areas of extensive tissue overgrowth. Indeed, growing evidence suggests PROS mutations cause overgrowth non-cell-autonomously, although mechanisms of this are poorly understood. Here, we develop mosaic zebrafish models of PROS with overexpression of orthologous hotspot *pik3ca* mutations (*pik3ca^PROS^*) to visualize and examine the effects of mutated cells on early development, in whole live animals. Reminiscent of PROS, we observe a spectrum of embryonic vasculature malformations (VMs), accompanied by larval muscle and bone overgrowth. Surprisingly, VMs only rarely expressed *pik3ca^PROS^* in constituent endothelial cells, with *pik3ca^PROS^*-expressing cells often closely abutting malformations instead. Single-cell transcriptomics of *pik3ca^PROS^*mosaic zebrafish prior to VM onset revealed that most *pik3ca^PROS^* cells are relatively immature and developmentally inert or constitute a small minority of mesodermal-derived cell types. Despite this constriction, global changes to cell fate were evident, alongside pervasive, pan-lineage abnormalities of gene expression, and rewiring of predicted ligand-receptor communication between lineages. We propose that targeting signals that indirectly propagate overgrowth through non-cell autonomous mechanisms - as well as PI3K activation itself - is worthy of therapeutic investigation.

**Significance statement:** Patchy, or mosaic, activating mutations in *PIK3CA* cause asymmetric human overgrowth due to aberrant hyperactivation of phosphoinositide 3-kinase (PI3K) signaling. Overgrowth prominently affects blood vessels and may be severely debilitating. While causation by *PIK3CA* mutations is clear, the explanation for the extent and pattern of associated overgrowth is not. Leveraging the power of zebrafish models for observation of early development, we now provide evidence for widely pervasive developmental effects of PIK3CA mutations extending beyond transcriptionally and phenotypically affected cells and lineages. This suggests potential therapeutic value of targeting secondary effects of PIK3CA activation as well as the activated PI3K itself.

## Introduction

Coordination of lineage determination and growth rates within and among cell lineages is central to healthy human development. Class 1 phosphoinositide 3-kinases (PI3Ks) are one crucial group of signaling enzymes that mediate such coordination. PI3Ks transduce numerous upstream growth cues into widely ramifying downstream effects on cell growth, metabolism and other behaviors (1). They do this in a cell type, developmental stage and tissue specific manner. These growth cues include hormones and growth factors which activate receptor tyrosine kinases. The dominant class 1A PI3K subtypes regulating cell growth and metabolism in non-hematopoietic cell lineages include the p110α catalytic subunit encoded by *PIK3CA*, acting in concert with one of several regulatory subunits (2, 3). The heterodimeric enzyme is typically referred to as PI3Kα.

Hyperactivation of PI3Kα is common in solid human cancers, most often attributable to somatic activating *PIK3CA* mutations. The mutations with the greatest hyperactivity fall in ‘hotspot’ regions at the C terminal of the kinase domain (Histidine 1047) or in the helical domain (Glutamates 542 or 545) (4). These same mutations, if acquired during human development, also cause a wide range of asymmetric overgrowth disorders. Many different disease labels are used for these based on phenotypic descriptors, and the term *PIK3CA*-related overgrowth spectrum (PROS) is now used as an umbrella term for the whole range, reflecting its shared etiopathogenesis (5, 6). PROS ranges from very localized overgrowth, for example a circumscribed vascular malformation or single digit overgrowth, to severe and sometimes life-threatening multisystem disorders such as CLOVES syndrome (Congenital Lipomatous Overgrowth, Vascular Malformations, Epidermal Nevus, Spinal/Skeletal anomalies/Scoliosis) (5, 7). Vascular complications such as thromboembolism, hemorrhage, mass effects or high output heart failure are major causes of morbidity and mortality in PROS.

The wide phenotypic spectrum of PROS has most commonly been rationalized in terms of the timing and cell of origin of the founder *PIK3CA* mutation, which determines the anatomical site(s) of overgrowth, with the *PIK3CA* genotype (hotspot *vs* non hotspot) an important accessory factor determining the rate of overgrowth of affected tissues. This formulation fails to provide a complete account of the observed phenotypic heterogeneity, however. One unexplained observation is that not all tissues are equally affected in PROS. Mesoderm-derived veins, capillaries, lymphatics, adipose tissue, skeletal muscle, and bone are commonly and severely affected, for example, as are ectoderm-derived brain and peripheral nerves. In contrast, mesoderm-derived arteries and hematopoietic cells, and endoderm-derived organs are rarely affected (4, 7). Given the presumed stochastic nature of spontaneous *PIK3CA* mutagenesis, this implies that mutations are excluded from, not tolerated in, or not phenotypically penetrant in some lineages (4).

Another important observation is that the burden of *PIK3CA* mutation is variable, but often in the range of <1-10% in tissue biopsies, even from macroscopically severely affected regions, meaning that the bulk of sampled cells are genetically wild-type (WT) (4, 8–10). This will partly reflect sampling of a genetically mosaic tissue, but the extent of this mismatch also suggests that *PIK3CA* mutations may exert both cell-autonomous and non-cell-autonomous effects on tissue growth. Work emerging using other model systems has shown that non-blood vessel cells, such as PIK3CA-mutant lymphatic endothelial cells or Schwann cells, can increase PI3K activation in neighboring endothelia to worsen vascular lesions and drive proliferation of WT cells, through recruitment of macrophages or secretion of pro-inflammatory eicosanoids and cytokines to drive proliferation of WT cells (11–14). Beyond its conceptual interest, identifying mediators of such non-cell-autonomous mechanisms may be of translational value if these secondary drivers of pathologic growth can be targeted. This is highly relevant in PROS as available potent PI3K inhibitors have limited efficacy in only a subset of patients in studies so far (15, 16), underscoring the need for better classification and understanding of the wider signaling landscape within VMs to ensure optimal treatment strategies (17).

Dissecting how the presence of *PIK3CA*-mutated cells corrupts normal development is ideally interrogated *in situ* in developing animals. Numerous animal models of PROS have indeed been reported to date, using strategies based on endogenous (18–20) or transgenic overexpression of *PIK3CA* hotspot variants (21–27), or a strongly activating but synthetic *Pik3ca* allele (28, 29). However, although such strong genetic *Pik3ca* activation has been shown to drive overgrowth in numerous lineages, the early events in overgrowth *in utero* are not visible in mammals. We now present a zebrafish model of PROS circumventing this limitation. The genetic tractability and transparency of zebrafish from their first cell divisions into adulthood means that critical stages in disease onset and evolution hidden *in utero* can be visualized and followed at cellular resolution from disease onset to adulthood in an intact organism (30). The large clutch size and rapid development of zebrafish also offer high experimental throughput to enable rapid testing of multiple PROS alleles in distinct cellular lineages.

We use this new model to investigate the cellular and transcriptional landscape at the onset of PROS vascular overgrowth. We show that overexpression of *pik3ca* hotspot alleles (henceforth collectively “*pik3ca^PROS^*”) causes extensive vascular malformations, despite the majority of *pik3ca^PROS^*cells localizing adjacent to, rather than within lesions. Overexpression limited to the mesoderm had widespread effects on subsequent development, manifesting as aberrant partitioning of mutant cells among mesoderm-derived lineages. Single cell transcriptomics suggested more pervasive effects on gene expression and signaling in non-mesodermal lineages, with potential loosening of the normal pattern of developmental signal crosstalk among key embryonic structures. These findings consolidate evidence for developmental effects of *pik3ca^PROS^* mosaicism that extend far beyond affected cells. This aligns with observations made in human PROS, presents a versatile animal model for further mechanistic interrogation, and opens new possibilities for accessory therapeutic targets beyond PI3K itself.

## Results

### Mosaic *pik3ca* hotspot transgenes cause vascular malformations in zebrafish embryos

Zebrafish Pik3ca protein has 91% homology with human PIK3CA, and AlphaFold predicts closely similar higher order structure to the experimentally determined structure of *PIK3CA*, evidenced by nearly identical positioning of PROS-related hotspot variants in both species **(Figure 1A)**. To develop a zebrafish *pik3ca-PROS* model, we generated Tol2-based vectors permitting *β-actin* promoter-driven (ubiquitous) expression of zebrafish *pik3ca*; either wild-type (WT) or harboring one of the hotspot mutations, namely *H1048R*, *H1048L* (orthologous to human *H1047R* and *H1047L* respectively), *E542K*, or *E545K* **(Figure 1B)**. A downstream IRES-*mScarlet* sequence enabled co-expression of the mScarlet reporter with mutant *pik3ca*. Additionally, the presence of an uncoupled secondary *cmcl2:GFP* cardiac reporter was included to provide a secondary readout of transgene integration. The *pik3ca^H1048R^-mScarlet* construct was introduced into WT embryos to produce F0 mosaic transgenic embryos, which were raised and screened for transgene integration and overgrowth.

**Figure 1:**
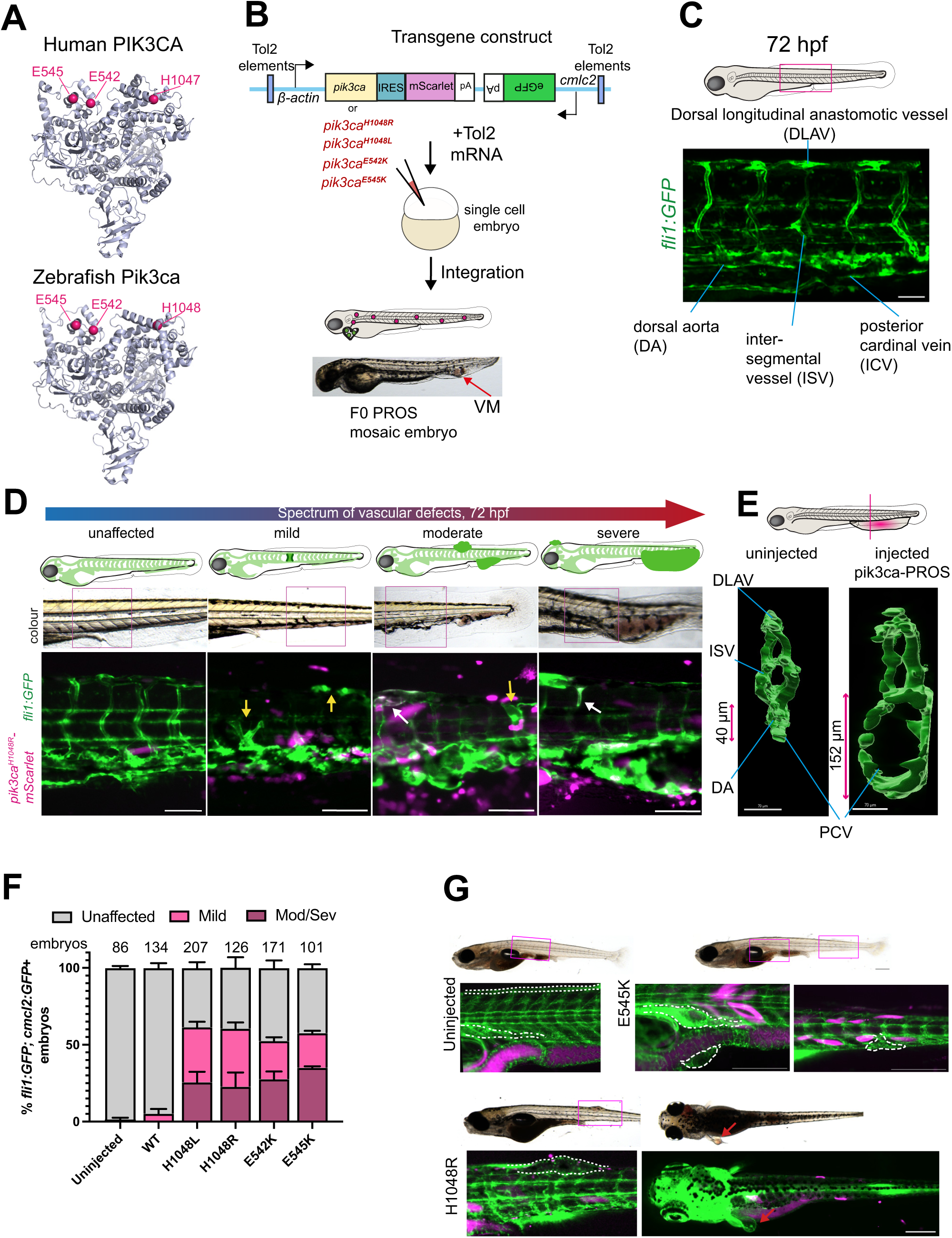
Mosaic *pik3ca* hotspot transgenes cause vascular malformations in zebrafish embryos. A) The structure of human PIK3CA and the predicted structure of the zebrafish Pik3ca orthologue is remarkably conserved, with one additional amino acid in zebrafish Pik3ca shifting the kinase domain amino acids by 1. The position of E542, E545 and H1047 residues found mutated in PROS (and the orthologous E542, E545 and H1048) are labelled in magenta and are predicted to occur in very similar structural regions of the protein in both human and fish. B) Schematic representation of the transgenic *pik3ca^PROS^* constructs generated for this study, containing *β*-*actin:pik3ca^PROS^*-*mScarlet* and the cardiac reporter *cmlc2:GFP* flanked by Tol2 response elements, and is co-injected with *Tol2* mRNA into 1-cell zebrafish embryos. Embryos were screened for successful integration of the transgene from 48 hpf onwards, indicated by mosaic cardiac *cmlc2:GFP*, and *mScarlet* expression. At 72 hpf, some PROS mosaic embryos had tail vascular malformations. C) Maximum Z-projection of a 72 hpf *Tg(fli1:eGFP);casper* tail region, showing typical endothelial vessel structures marked by *fli1:eGFP* fluorescence (green). Scale bar = 50 μm D) Representative diagrams and fluorescent microscopy images of the spectrum of vascular phenotypes seen in *Tg(fli1:eGFP)* embryos injected with *pik3ca^H1048R^-mScarlet*. These ranged from unaffected and mild, to moderate and severe. *Fli1:eGFP* expression in unaffected embryos is as uninjected. Embryos scored ‘mild’ displayed expansion and mispatterning of isolated inter somitic vessels that was only visible when viewing eGFP fluorescence. Moderate and severe scores were applied to embryos where blood pooling was evident under brightfield, particularly obvious in the PCV. Yellow arrows indicate mis-patterned vessels in ‘mild’ embryos, and white arrows indicate overlap of *fli1:eGFP* (green) and *mScarlet* (magenta) expression in moderate and severe examples. Quantification of this and numbers screened can be seen in Figure 3. Scale bars = 100 μm. E) Cross sections of trunk blood vessels at 72 hpf, showing the dramatic expansion of the PCV. F) Quantification of these vascular malformation phenotypes for each construct, scored based on severity, as a percentage of the total numbers injected. Numbers indicate *fli1:eGFP; cmcl2:GFP+* embryos screened for vessel malformations at 72 hpf, representative of three biological repeats per construct. Error bars represent S.E.M. G) Representative images of juvenile *Tg(β-actin:pik3ca^E545K^-mScarlet; fli1:eGFP); casper* (top) and *Tg(β-actin:pik3ca^H1048R^-mScarlet; fli1:eGFP* fish that harboured no detectable vascular anomalies at 72 hpf, but were found to have vascular anomalies when imaged two weeks later (*E545K*-injected – 6/16, *H1048R*-injected – 3/9). Dotted lines encompass vessel malformations in PROS animals, and equivalent regions in control

Many *pik3ca^H1048R^-mScarlet-*injected embryos appeared morphologically normal, but 15% developed vascular lesions, observed as blood pooling and poor circulation most evident at 72 hours post-fertilization (hpf) **(Figure 1B)**. To investigate this phenotype further, we repeated *pik3ca^H1048R^-mScarlet* transgene injections into the *Tg(fli1:eGFP)* endothelial reporter line, enabling co-visualization of blood vessels and transgene expression (31). *Fli1:eGFP* is first detectable during somitogenesis, during which angioblasts migrate from the lateral plate mesoderm (LPM) between the 10 and 18 somite stage (∼16 hpf) towards the midline, with the main embryonic artery and vein being formed at 24 hpf, and trunk circulation being completed at 48 hpf (31, 32). In the trunk of zebrafish embryos at 72 hpf, arteries and veins are well established, with regularly-spaced intersegmental vessels (ISVs) connecting the ventral major aortic and venous vessels with the dorsal longitudinal anastomotic vessel (DLAV) (31, 33) **(Figure 1C)**. Consistent with the variability observed in PROS, the severity of vascular malformations resulting from transgene injection ranged from mild to severe, the latter obvious from dramatic enlargement of the major blood vessels, causing blood pooling in the tail or head, with some vascular overgrowths large enough to compromise swimming (**Figure 1D**).

In the same system, we next tested the propensity of other PROS hotspot variants to cause vascular malformations, injecting WT, *H1048R, H1048L, E542K* or *E545K pik3ca^PROS^* constructs into *Tg*(*fli1:eGFP)* embryos. Transgenic expression of *pik3ca^WT^-mScarlet* did not induce overgrowth (**Supplementary Figure 1A**). However, all four hotspot variants, whether affecting the kinase (*H1048R/L*) or helical domains (*E542K/E545K*) of Pik3ca, reproducibly caused a spectrum of vascular anomalies, from mild mis-patterning of *fli1:eGFP+* ISVs only evident by fluorescent transgene expression pattern, to severe balloon-like overgrowth, of the major vessels in the tail **(Figure 1D,E Supplementary Figure 1A)**. Severity scoring of malformations for each of the four PROS alleles and of WT or uninjected controls showed that transgenic expression of mosaic *pik3ca^PROS^*variants in zebrafish caused vascular malformations at similar levels of severity (**Figure 1F**).

To test whether our model could recapitulate other overgrowth elements seen in PROS, we selected 72 hpf embryos displaying a high level of *pik3ca^PROS^*transgene integration but appearing otherwise normal, and grew them to larvae stages (to 14 days). Transgene integration was assessed either by high numbers of *mScarlet+* cells or by the surrogate marker of expression of the co-injected heart-targeted *cmcl2:GFP* reporter in most cardiac cells. We imaged these fish as they developed into larvae. Despite no visible vascular abnormality in the embryos, these fish still developed vascular malformations by larval stages, seen as thickening of vessels, or as vascular protrusions from the skin **(Figure 1G)**. Aside from vascular malformations, some *pik3ca^PROS^* zebrafish larvae also showed appreciable *mScarlet+* expression and overgrowth in the musculature, as well as premature mineralization in the notochord, fusing and thickening of vertebrae, and abnormal patterning of hypural cartilage in the tail **(Supplementary Figure 1B-D)**. These findings suggest that in the fish model, as in human PROS, the trajectory of malformations is not exclusively determined in embryos, and overgrowth can sometimes manifest or progress post-embryonically, notably in tissues of mesodermal origin.

### Evidence for cell-autonomous and non-cell-autonomous mechanisms of *pik3ca^PROS^* vascular overgrowth

Surprisingly, we observed that only a minority of endothelial cells within vascular malformations expressed both mScarlet and eGFP fluorophores, proxies for *pik3ca^PROS^* and *fli1* expression, respectively. We used confocal microscopy to build a 3D image of the vascular lesions in higher resolution to investigate this further **(Figure 2A; Movie 1)**. In one example of a mild VM, we observed thickening of the DLAV without mScarlet cells in the lesion itself, but with *pik3ca-PROS*-*mScarlet+* cells abutting the overgrown region **(Figure 2A; Movie 2)**. In other moderate and severe vascular malformations, *pik3ca^PROS^-mScarlet+* cells were again most often found adjacent to malformations rather than comprising abnormal vessel walls (**Figure 2A; Movie 3**). Indeed, GFP and mScarlet double positive cells accounted for <5% of total lesion volume on average **(Figure 2B).** This provides circumstantial evidence for the hypothesized non-cell autonomous mechanisms in PROS. Specifically, it argues that unidentified *pik3ca^PROS^* cells in or close to the vascular niche may exert non-cell-autonomous effects upon WT endothelial cells.

**Figure 2:**
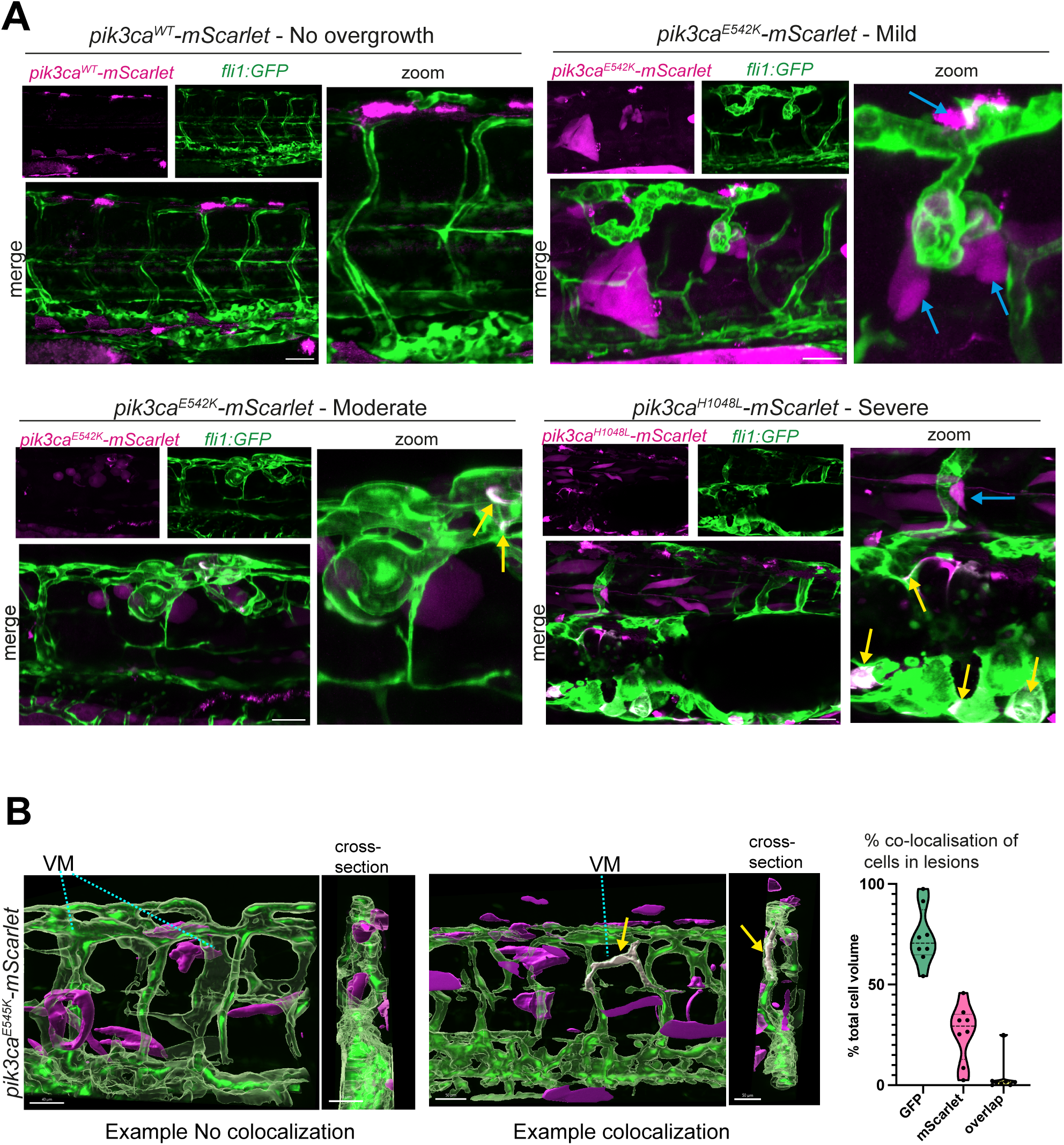
Evidence for cell-autonomous and non-cell-autonomous mechanisms of *pik3ca^PROS^* vascular overgrowth. A) Representative maximum projection confocal images of 72 hpf *Tg(fli1:eGFP);casper* embryos expressing either wildtype *pik3ca-mScarlet* (no overgrowth) or a *pik3ca^PROS^* hotspot variant and exhibiting a mild, moderate or severe vascular phenotype. For each, separated *mScarlet* fluorescence (magenta), *fli1:eGFP* channels (green) and merged channels are shown, with a zoomed area to the right. Blue arrows indicate *mScarlet* cells in close proximity to deformed vessels. Yellow arrows indicate white cells co-expressing *fli1:eGFP* and *mScarlet*. Scale bars = 50 μm B) Quantification of *mScarlet* cells by volume within GFP+ endothelial cells. Imaris was used to visualize confocal Z stacks of vessel overgrowth at 72 hpf. 3D structures of each channel, plus a co-localization channel were converted into surfaces, enabling volume calculations for vessels (green), mScarlet+ cells outside vessels (purple) or areas of colocalization (white). Volumes of each were expressed as percentage of total cell volume in 20x magnification view, dots indicate individual images (8 lesions were analysed in total).

### Early mesodermal *pik3ca^PROS^* transgene expression causes endothelial mis-patterning

Many tissues prominently affected in PROS are mesoderm-derived, indicating biases in cell lineage contribution to overgrowth, and the possibility of spatial and temporal windows of especial vulnerability to PROS mutations during development (4). We therefore sought to investigate the consequences of disease onset in the early mesoderm, which has hitherto not been possible to observe *in vivo* in mammalian systems, by swapping our ubiquitous *β-actin* promoter for that of the pan-mesodermal marker *tbxta* (**Figure 3A, B**). The activation of *tbxta* prior to endothelial specification would also address the possibility that *mScarlet+* endothelial cells had been present during vasculogenesis but had either silenced transgene expression or died prior to imaging and scoring at 72 hpf. As all tested *pik3ca^PROS^* alleles gave rise to VMs with the same frequency and severity (**Figure 1F**), we focused on one hotspot variant for subsequent work, *pik3ca^E545K^,* for studying mesodermal PI3K gain-of-function.

**Figure 3:**
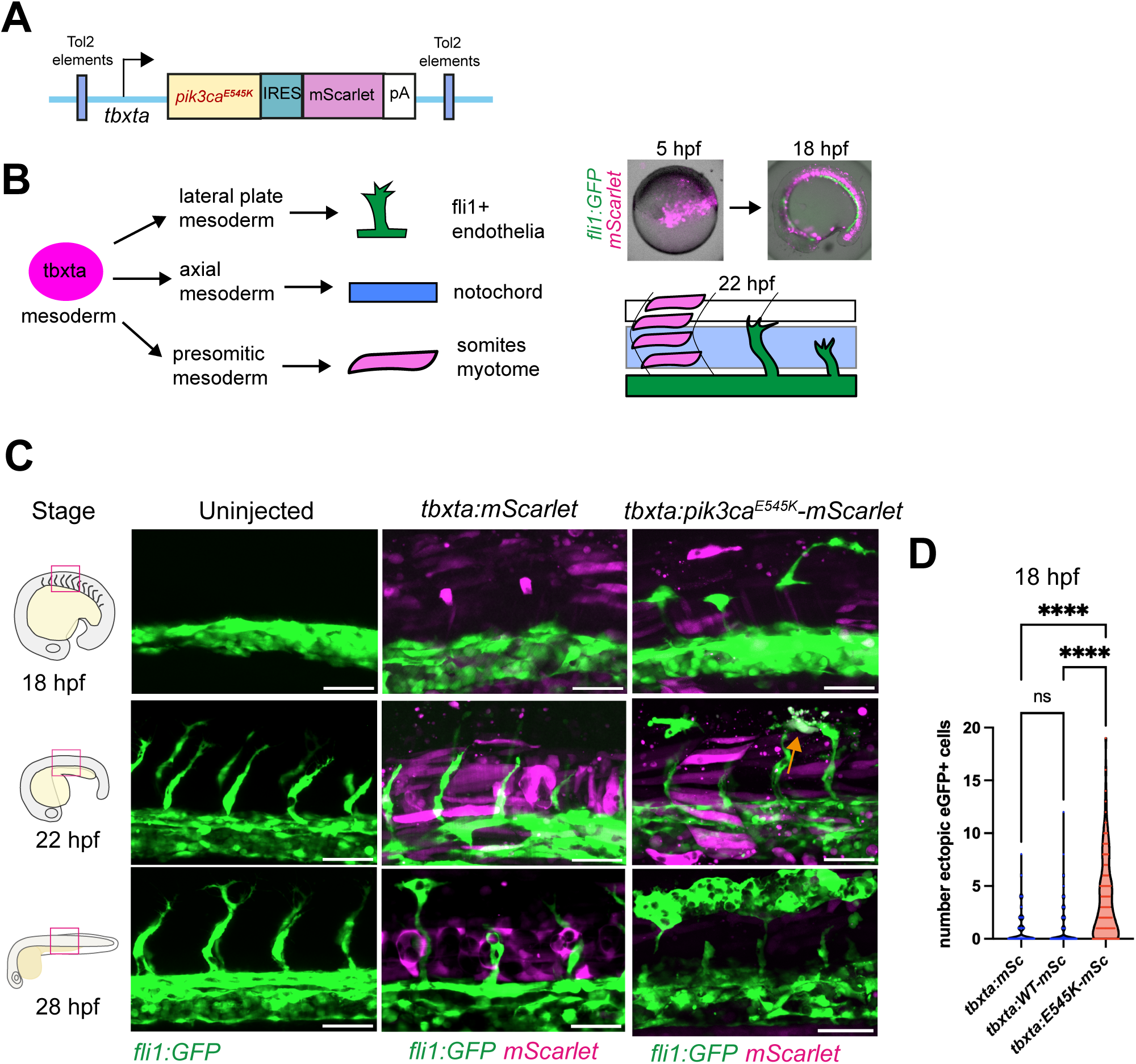
Early mesodermal *pik3ca^PROS^*transgene expression causes endothelial mis-patterning. A) Plasmid design of the transgenic construct *tbxta:pik3ca^E545K^-mScarlet*. B) Simplified diagram of a *tbxta* progenitor cell which may differentiate into the lateral plate mesoderm, the notochord, or presomitic mesoderm which gives rise to endothelia, the notochord or muscle respectively. Mosaic *tbxta:mScarlet* is visible at 5 hpf (representative images) before onset of *fli1:eGFP* reporter expression. By 18 hpf, *tbxta* is most highly expressed in the notochord. C) Representative maximum projection confocal images of embryos from a range of stages between somitogenesis (∼18 hpf) and tailbud (∼28 hpf) to visualise the start of *fli1:eGFP* expression and endothelial development. In uninjected embryos, lateral stripes of *fli1:eGFP* are present from 16hpf, with sprouting of ISVs occurring from anterior to posterior from around 19 hpf onwards. In *tbxta:pik3ca^E545K^-mScarlet* injected embryos, ectopic *fli1:eGFP* cells can be seen dorsal to the initial lateral plate mesoderm stripe. Scale bars = 50 μm D) Quantification of ectopic *fli1:eGFP+* cells in injected *tbxta:pik3ca^E545K^-mScarlet,* or injected with a similar construct lacking the *pik3ca-ires* element (unattached to the lateral mesoderm stripes of *fli1:eGFP*). 3 biological replicates shown with each dot representing 1 embryo. Significance was calculated with a two-tailed Mann Whitney test. ****=p<0.001.

*pik3ca^E545K^-mScarlet+* expressed from the *tbxta* promoter was as effective as the *β-actin* construct in inducing the *pik3ca^PROS^*vascular phenotype in *Tg(fli1:eGFP)* embryos (**Supplementary Figure 2**). *tbxta:pik3ca^PROS^-mScarlet* expression was visible from early gastrulation, before the earliest plausible stage of vascular malformation onset, as angioblasts migrate from the lateral plate mesoderm (LPM) towards the midline during somitogenesis (10-22 hpf) (31). At 18 hpf, *mScarlet* expression in injected controls was predominantly in the notochord and myotome, but co-expressed with very few GFP+ endothelial cells, as expected from known expression patterns of *tbxta* at this developmental stage (34) (**Figure 3C**). In controls, very occasional ectopic *fli1:eGFP* cells were observed dorsal to the developing DA and unattached to the nascent vasculature, thus seemingly independent of normal ISV sprouting, which occurs from 20 hpf (**Figure 3C, Movie 4**). In embryos overexpressing *pik3ca^PROS^* however, the frequency of such ectopic *fli1:eGFP* cells was markedly increased (**Figure 3D**). These ectopic cells persisted through ISV sprouting and subsequent branching to form the DLAV at ∼22 hpf. Occasionally, at this time point, we observed rare examples of *eGFP* and mScarlet double positive cells dying (**Figure 3C**, orange arrow). By 28 hpf, we observed DLAV cells forming premature dilated vessels (**Movie 5**).

We thus demonstrate that *pik3ca^PROS^* expression in mesoderm alone disrupts the earliest stages of vascular development, causing dysregulation of endothelial cell sprouting and branching. Occasional *mScarlet*-expressing endothelial cells were present, some of which were observed dying during aberrant development, which may indicate heightened sensitivity of endothelial cells to high levels of *pik3ca^PROS^* transgene overexpression. However, as in later vascular development, vascular defects in this model appear to be largely a non-cell autonomous effect of *pik3ca^PROS^*expression on *fli1:GFP+, mScarlet+* endothelial cells.

### Mosaic mesodermal *pik3ca^PROS^* expression induces pan-lineage dysregulation

Given this evidence for non-cell autonomous effects of mesodermal *pik3ca^PROS^* expression on early vascular development, we next sought to interrogate more agnostically the extent of the developmental disruption. We used FACS to sort live cells from *tbxta:pik3ca^PROS^-mScarlet* and *tbxta:mScarlet* injected controls at 19 hpf and subjected these to single-cell RNA sequencing (scRNA-seq) (**Figure 4A**). Using Seurat, we identified 40 distinct cell populations common to ‘control’ and ‘PROS mosaic’ embryos. These were named using a combination of marker gene expression patterns and projection of two previously published scRNA-seq datasets onto the data (**Figure 4B, Supplementary Figure 3A, B**) (35, 36). This confirmed the presence of all cell lineages expected at this developmental stage, including endothelial cells, CNS, epithelial, lateral plate and presomitic mesoderm (LPM, PSM), and notochord (**Figure 4B, C**).

**Figure 4.**
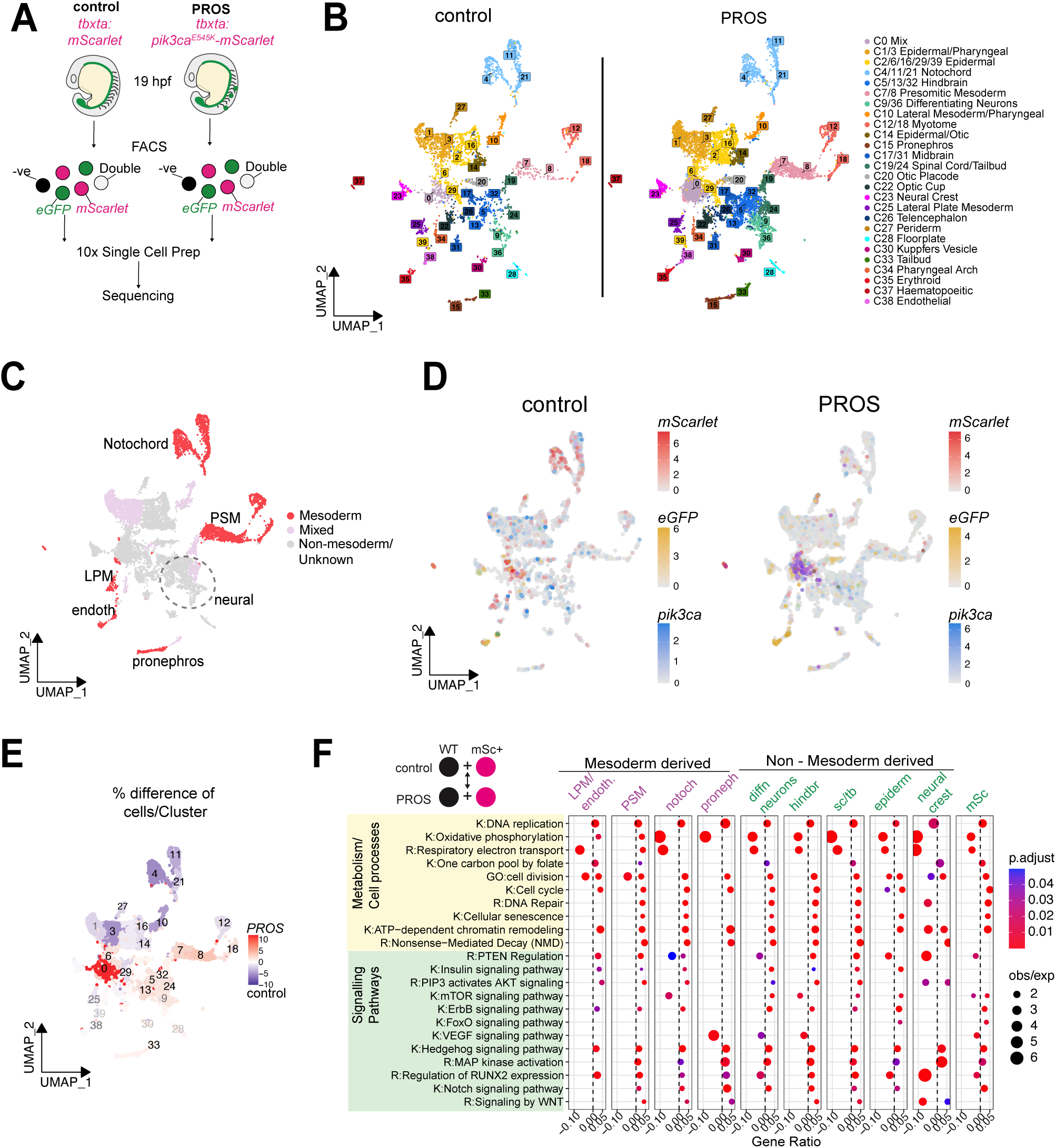
Mosaic mesodermal *pik3ca^PROS^* expression induces pan-lineage dysregulation. A) Schematic of experimental workflow for scRNA-seq. *Tg(fli1:eGFP)* embryos were injected with *tbxta:mScarlet* (control) or *tbxta:pik3ca^E545K^-mScarlet* (PROS) and grown until 19 hpf, dissociated, and all live cells sorted by FACS before processing for 10x sequencing. B) UMAP showing 40 cell clusters found the aggregated dataset, split by experimental condition. C) UMAP of aggregated dataset coloured by mesodermal or other germ layer origin D) FeaturePlots split by experimental condition, showing expression levels of *mScarlet* (red scale)*, eGFP* (yellow scale) and *pik3ca* (blue scale) E) The numbers of cells per cluster was expressed as a percentage of total cells for each experimental condition, and then the difference between percentages in PROS and control clusters visualised in a UMAP heat plot. Blue indicates a higher percentage of control cells relative to PROS, and red indicates a higher proportion of PROS relative to control cells. F) Deseq2 DE analysis and gene enrichment analyses using GO (G:), KEGG (K:) and Reactome (R:) databases were performed to compare PROS mosaic vs control cells in various cluster groups, or in *mScarlet+* cells across the dataset.

One clear difference between control and *pik3ca^PROS^* embryos lay in so-called cluster ‘0’, which could not be mapped to a uniquely annotated lineage, instead featuring weak signatures of many different cell types, hinting at a “confused” lineage identity (**Supplementary Figure 3C**). We confirmed upregulation of the *pik3ca* transcript itself in this cluster, largely co-segregating with *mScarlet* in PROS mosaic animals (**Figure 4D**). Velocity analysis showed a lack of clear developmental directionality of transcription in PROS mosaic embryos compared to the small corresponding cluster in control animals. In contrast, velocity in the PSM and neural lineages was increased, suggesting non-cell autonomous perturbations of lineage transitions in both mesodermal (*tbxta*-driven) and non-mesodermal (i.e. neural) lineages (**Supplementary Figure 3D**).

The changes in relative sizes of different cell clusters between experimental conditions was assessed by expressing the number of cells within each cluster as a percentage of the total cell number for control and PROS datasets, respectively. After calculating the differences in these percentages, a dramatic expansion of cluster 0 was observed in PROS mosaic embryos (**Figure 4E; Supplementary Figure 3E**). Some clusters of mesodermal origin also showed expansion in PROS mosaics, as may be expected from mesodermal activation of PI3K signaling, but, surprisingly, this was not true of all mesoderm-derived clusters, and ectoderm-derived neural clusters were also enriched in PROS mosaics (**Figure 4D, E; Supplementary Figure 3E**). This suggests that *pik3ca^PROS^* cells have developmental effects extending beyond mesodermal lineages themselves, unbalancing cell fate decisions and thereby cell type proportions in the embryo.

In PROS mosaic embryos, 319 genes were significantly upregulated in mScarlet+ PROS cells relative to wildtype in PROS mosaics, including, as expected, *pik3ca, tbxta* and *mScarlet*, but fewer were upregulated in PROS mosaics compared to control embryos (**Supplementary Figure 4A**). Analysis of curated transcriptomic “footprints” of pathway activity indicated that PI3K-coupled programs were significantly enriched in *mScarlet*+ cells in PROS mosaics relative to their WT *mScarlet-* counterparts, indicating increased PI3K activity as well as *pik3ca^PROS^* overexpression in transgenic cells (**Supplementary Figure 4B**). Interestingly, the same analysis revealed enrichment for signatures of apoptosis, DNA damage repair, G2/M checkpoint activation and p53 activity, suggesting that *pik3ca^PROS^* expression may ultimately be toxic. Such negative selection of *pik3ca^PROS^* has previously been invoked as a possible explanation for the relative lack of PIK3CA mutations in some human cell lineages. Surprisingly, few genes were significantly upregulated in PROS mosaic cluster 0 relative to control, although footprint analysis did suggest significant enrichment of MYC-Targets and Apical Junction components (**Supplementary Figure 4C**), although the uncertain nature of cluster 0 in control embryos complicates interpretation of this comparison.

In other cluster groups, we found widespread enrichment of Kegg or Reactome pathway components directly related to PI3K signaling in clusters from the PROS zebrafish, including modules related to Pten, Insulin, PIP3, mTor and Foxo1 (**Figure 4F**). Other gene sets were also frequently enriched, including Notch, Wnt and Mapk pathways, as well as shared responses to cell metabolism and cell division processes. Strikingly, these transcriptional changes extended to clusters in the PROS zebrafish of non-mesoderm origin that expressed almost no mScarlet cells (**Figure 4F**). Moreover, these gene expression patterns persisted when we removed the PROS *mScarlet+* cells from the analysis entirely, indicating that otherwise wild type mesodermal and non-mesodermal lineages are responding transcriptionally to PROS cells even without expressing *pik3ca^PROS^* themselves (**Supplementary Figure 4D**).

### *pik3ca^PROS^* elicits widespread transcriptional changes in ligand-receptor expression patterns

Finally, we sought evidence from single cell transcriptomic data implicating potential mediators of the observed non-cell autonomous effects of genetic PI3K hyperactivation. We did this using CellChat and NICHES as analytic tools to infer communication among cell lineages using the proxy of expression patterns of ligand/receptor pairs (37, 38). Analyses of the top 10% predicted cluster interactions by CellChat suggested major transcriptional changes in signalling mediators in PROS embryos. In WT embryos the transcriptomic landscape appeared dominated by reciprocal expression of ligand receptor pairs (loosely “signaling traffic”) among the notochord, and mixed epithelial/pharyngeal lineages, while in PROS embryos a more distributed pattern was seen, with presomitic mesoderm and various neural lineages much more prominent (**Figure 5A**).

**Figure 5.**
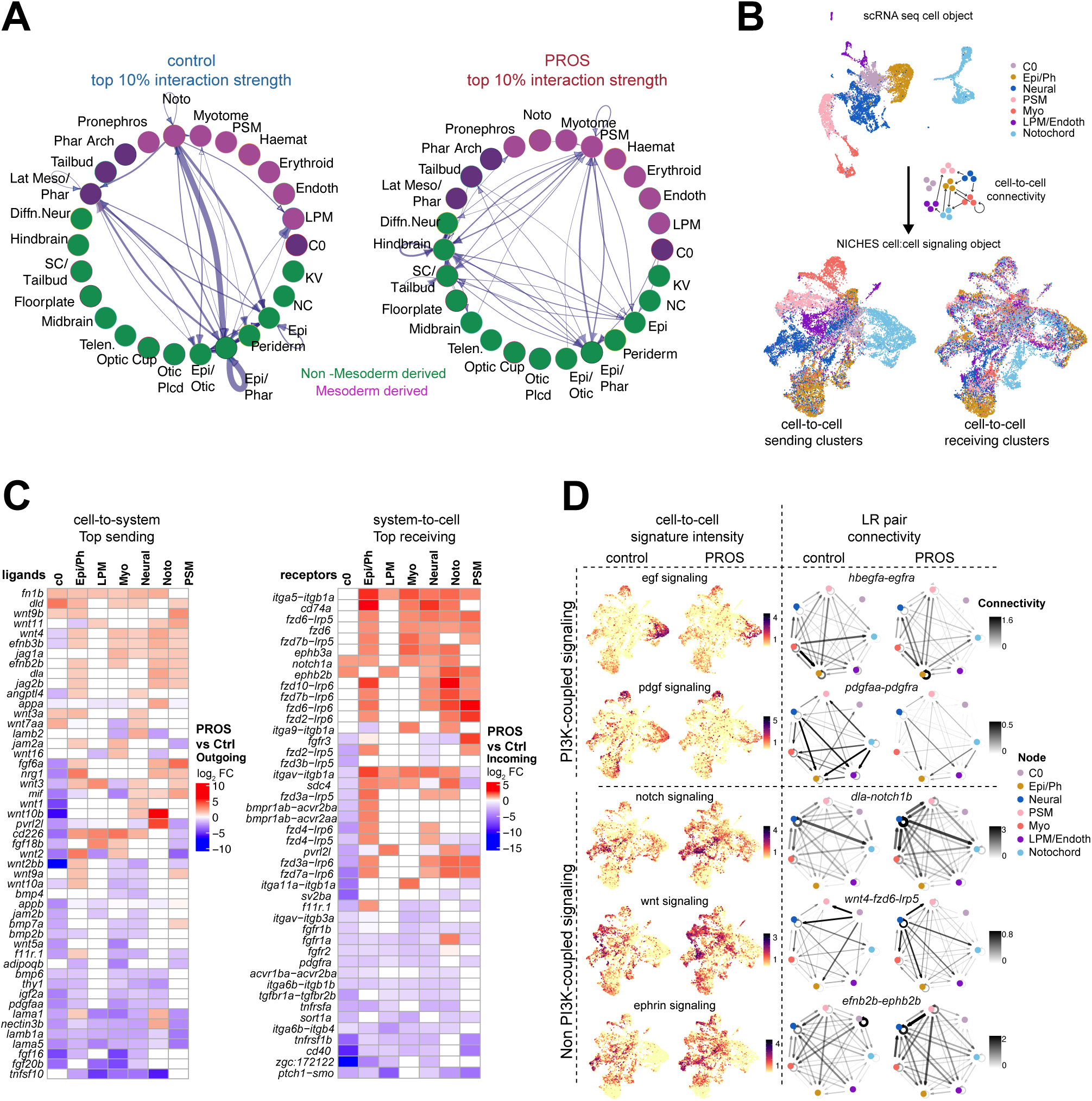
*pik3ca^PROS^*promotes widespread transcriptional changes in ligand-receptor expression patterns. A) CellChat was used to identify potential interactions between ligand and receptor pairs in clusters. The top 10% strongest pathway interactions of ‘control’ cells relative to ‘PROS’ and vice versa shown, normalised to cell number. Blue arrows indicating whether a cluster is a sender or a receiver of the signal, with arrow thickness proportional to connection strength. Abbreviations: LPM=lateral plate mesoderm, PSM, presomitic mesoderm, Noto = notochord, Endoth=endothelial, KV = kuppfer’s vesicle, NC=neural crest, Epi=epiderm, Epi/Phar epidermal/pharyngeal, Otic Plcd=otic placode, Telen=telencephalon, SC=spinal cord, Diffn. Neuron = differentiating neurons, B) UMAP showing merged data to illustrate the methodology of NICHES, whereby a subset of clusters from Figure 4 is reclustered (Top UMAP). Ligand-receptor interactions between each group were quantified using NICHES, converting it into a signaling object of cell:cell interactions (Bottom UMAPs, grouped by either sending cluster type (left) or receiving cluster type (right). Colours in interaction maps were retained from the original clusters in the top UMAP. Epi/Ph = epiderm/pharyngeal mix, PSM = presomitic mesoderm, Myo=myotome, LPM/Endothelial = grouped lateral plate mesoderm and endothelial clusters. C) Heatmap depicting log_2_ fold change in expression of the top differentially expressed “sending” ligand (left) and “receiving” receptor (right) genes in PROS relative to control embryos, clustered by row. Ligand-receptor mechanisms were aggregated by common ligand or common receptor (unaggregated data are shown in **Supplemental Figure 4C**). Only differences for which p.adj<0.05 by Wilcoxon test between PROS and WT are shown. D) Examples of pathways showing dysregulated expression of ligand-receptor (LR) pairs. The expression values of LR pairs for each pathway were averaged as a module score. FeaturePlots (left) show intensity of this module for each cell:cell interaction, split by experimental condition. Examples of circus plots for each pathway are shown on the right. Signal direction between clusters (nodes) is shown by arrows, with darker and thicker arrows corresponding to greater connectivity strength. Connectivity scale intensity is matched between control and PROS plots for each LR pair.

To assess potential specific signaling pathways driving these predicted changes, we used the NICHES algorithm, which analyses ligand-receptor mRNA expression in one-to-one pairs of cells to infer communication (38). We subsetted our data to include the clusters found most active by CellChat and ran NICHES to convert the single-cell gene expression dataset to a single-cell signalling dataset based on LR signaling pattern genes, grouped by sending clusters, receiving clusters, or individual pair-wise interactions (**Figure 5B)**. To identify potential changes in cell signalling capacity between lineages in PROS mosaic animals relative to control, we performed DE analysis on ‘sending’ and ‘receiving’ transcriptional signatures for each cluster (**Supplementary Figure 4E**). To gain a comprehensive overview of signaling changes, we simplified the outputs by grouping and averaging ligand and receptor fold changes respectively (**Figure 5C**). We observed a marked increase in Notch, Wnt and Ephrin ligand-receptor pairs in both mesodermal and non-mesodermal lineages, but again cluster 0 exhibited general downregulation of sending and receiving signaling components. This is in keeping with the non-cell autonomous responses to small numbers of *pik3ca^PROS^*cells in the PROS zebrafish, and the relatively inert state of C0 (**Figure 5C**; **Supplementary Figure 3 C,D**).

We next calculated ‘module scores’ by averaging gene expression of all ligand-receptor pairs within a pathway (e.g. all combinations of *jag*/*dll*/*dld*-*notch1a/1b/2/3* for notch signalling) and compared the intensities of this score on cell-to-cell UMAPs, split by experimental condition (**Figure 5D)**. Comparing the connectivity of PROS mosaics to control, we observed a reduction in extent and intensity of mRNA expression of *egf* and *pdgf* ligand-receptor pairs in PROS, further highlighting disruption of normal signaling through Pik3ca throughout the embryo (**Figure 4F, 5D**). In contrast, we observed widespread upregulation of pairs of *notch*, *wnt* and *ephrin* signaling genes (**Figure 5D**). Focusing on individual ligand-receptor pairs, we found evidence for transcriptional “rewiring” of these pathways between lineages. For example, *dla-notch3, wnt4-fzd5-lrp6* and *efnb3b-epha4a* ligand-receptor pair expression is increased for neural, PSM, myotome and Epi/Ph nodes (**Figure 5D**).

We conceptualize these data to indicate that *pik3ca^PROS^*has significant effects on cell fate, behavior and likely signaling which ramify across many lineages, indicating pervasive and disruptive non-cell-autonomous effects on early development. Overall, these analyses highlight transcriptional changes in ligand and receptor gene expression that may contribute to the non-cell-autonomous phenotypes observed and serve as potential drug targets worthy of further investigation.

## Discussion

Over ten years since mosaic, activating PIK3CA mutations were identified as the cause of a group of asymmetric overgrowth disorders (39, 40) much has been learned. The wide human clinical spectrum has been delineated, the natural history described, and nomenclature harmonized (5, 6). Furthermore, case series and registry based open label studies have established that the PI3Kα inhibitor alpelisib is of some clinical utility, leading to licensing in the USA (16, 29) and establishment of an international randomized clinical trial still underway (NCT04589650) (41). However, despite this major progress, important fundamental and translational questions about PROS remain. Specifically, the explanation for the skewed appearance of PROS-related pathology in some tissues but not others, and the involvement and extent of non-cell-autonomous mechanisms of overgrowth in PROS pathogenesis have eluded comprehensive analysis. The latter issue is of translational importance as it may yield novel diagnostic and therapeutic targets; given current evidence that alpelisib has only partial efficacy in only some patients with PROS (15), this is of great value.

Fundamental to translational progress is the fidelity of models to human pathology. A wealth of murine models has advanced our understanding of PROS, by showing definitive causal links between acquisition of PROS mutations and vascular malformations (22, 42), identifying critical time windows during which endothelia are most sensitive to excessive PI3K signaling (25), and demonstrating that PI3K and related pathway inhibition can mitigate or rescue important disease manifestations (14, 21–23, 42, 43). More recently, mouse PROS or cancer models have implicated signaling from non-endothelial cell types as drivers of vascular overgrowth (11, 25, 27), but otherwise few animal studies have focused on non-cell-autonomous contributors to PROS pathology to build a holistic view of multi-lineage interactions during early development.

Overexpression of an activating PIK3CA allele has been shown to recapitulate several aspects of PROS-related overgrowth in zebrafish (28), and the successful usage of zebrafish as a preclinical model of lymphatic overgrowth or neurovascular malformations (44, 45) encouraged us to develop patient-led zebrafish models to interrogate the complex interplay of Pik3ca genotype and lineage context, and to examine overgrowth dynamics at early stages of development. We now present a zebrafish model with mosaic overexpression of naturally occurring hotspot *pik3ca^PROS^* variants. This enables observation of the effects of PROS mutations *in vivo*, from onset, at cellular resolution for the first time. Our model recapitulates a spectrum of overgrowth severity affecting different cell lineages, as in PROS, with vasculature most prominently affected, while scRNA-seq suggests that *pik3ca^PROS^* expressing cells appeared ‘stuck’ in a less differentiated cell state. This is reminiscent of iPSC models of *PIK3CA^H1047R^* activation, where homozygous mutant cells show transcriptional remodeling and impaired differentiation (46). Our model also allows comparison of four different *pik3ca* hotspot gain-of-function variants in parallel, which proved to cause vascular overgrowth with similar frequency and severity. This argues against major differences in phenotypic expression across different hotspot Pik3ca variants, whether affecting helical or kinase domains, although the high level of expression in our model leaves open the possibility that differences may be unmasked at endogenous expression levels.

The non-cell autonomous influences of *pik3ca^PROS^* cells found in our *pik3ca^PROS^* model is in keeping with mounting evidence that paracrine signals from *PI3KCA^PROS^* cells corrupt neighboring wildtype cells. For example, eicosanoids secreted from *PIK3CA* mutant breast cancer cells induce proliferation in WT cells (13), paracrine VEGF and IL-6 signaling from mammary duct culture lead to dysfunction in neighboring innervating vasculature *in vitro* (12). Moreover, *in vivo* activating Akt mutations in murine mammary tissue cause hyperplasia of nearby WT cells in a paracrine manner (47), and lymphatic capillary endothelial cells expressing *Pik3ca^H1047R^* recruit VEGFC-producing macrophages which cause immune activation and worsening of vascular malformations in a mouse lymphoma model (11). Lastly, driving *Pik3ca^H1047R^*in murine pericyte progenitors induced widespread capillary and venous malformations as well as hypertrophy of Cre-targeted connective tissues (48). Paracrine signals also have a role in normal endothelial and hematopoietic development, as a subtype of endothelial cells derived from the paraxial mesoderm have been shown to support induction of hematopoietic cells from neighboring hemogenic endothelial cells in zebrafish (49). It is important to note that cell-autonomous and non-cell-autonomous contributions to VMs in our model are not mutually exclusive, and we did consistently observe minorities of *fli1:eGFP*+ endothelia co-expressing *mScarlet* indicative of a mixture of signaling mechanisms occurring in VMs. Likewise, gain-of-function *PIK3CA* mutations are found through genotyping endothelial cells in PROS patients, and are thought to be a driver of overgrowth in their wildtype counterparts. Therefore, both cell-autonomous and non-cell-autonomous signals likely underpin PROS pathology.

A limitation of our model, as for many other published models, is that it relies on transgenic *pik3ca* overexpression, although the variants we overexpressed were orthologues to human disease-causing mutations. The extent of overactivation through transgenesis is likely higher than seen in PROS where *PIK3CA* is regulated endogenously, meaning the lack of *mScarlet* in cell types most sensitive to PI3K levels, such as the endothelia, may be due to an intolerably high levels of PI3K signaling causing cell death, as we observed in some co-expressing cells, and enrichment of apoptotic transcriptional signatures in *mScarlet+* cells in PROS mosaic embryos. Furthermore, the timing of p110α activation is important for phenotype severity, with prenatal but not postnatal induction of *Pik3ca^E545K^* causing increased brain size, and timed induction of *Pik3ca^H1047R^* in endothelial cells revealing critical windows of PI3K sensitivity during development (25, 26). Further temporal resolution of *pik3ca^PROS^* during zebrafish development, and to an extent the ‘dosage’ of PI3K activation would be possible using a drug-inducible version of our transgene. Finally, further testing of our transgenic construct in older fish, perhaps with more restricted spatial expression of PROS mutations, is required to see if adipose tissue and other tissue defects observed in PROS occur, and to the same penetrance.

Our novel zebrafish *pik3ca^PROS^* model has enabled *in vivo* observation of early stages of evolution of PROS-like vascular malformations at a cellular level. It has moreover offered evidence for surprisingly pervasive developmental effects of mosaic *pik3ca* activation, including extensive non-cell autonomous effects ramifying beyond the affected lineage. This complements and extends existing models of PROS and provides a tractable platform for more in-depth interrogation of specific candidate mediators of such remote effects of *pik3ca* activation. Therapeutic targeting of these may in principle be a valuable adjunct or alternative to PI3K inhibition in future.

## Materials and Methods

### Structural comparison of zebrafish and human PIK3CA

Protein structures for human PIK3CA (AF-P27986-F1-v4) and the predicted structure of zebrafish Pik3ca (AF-F1QAD7-F1-v4) were visualized and annotated using PyMOL (Schrodinger).

### Fish husbandry, fish lines

Zebrafish were maintained in accordance with UK Home Office regulations, UK Animals (Scientific Procedures) Act 1986, amended in 2013, and European Directive 2010/63/EU under project licenses P8F7F7E52 and PP7317786. All experiments were approved by the Home Office and AWERB (University of Edinburgh Ethics Committee). Fish stocks used were wild-type AB, *Tg(fli1:eGFP*) (50), *casper* (51). Combined transgenic lines were generated by crossing. Adult fish were maintained at ∼28.5°C under 14:10 light-dark cycles. Embryos were kept at 28.5°C and staged according to the reference table provided by (52) and (53).

### *Pik3ca^PROS^* transgene plasmid construction

First, the *eGFP* sequence in the p3E Gateway entry vector IRES-eGFP-pA (54) was swapped for the *mScarlet* CDS (Thermo Fisher Scientific) with the HiFi DNA assembly kit (NEB) to make the p3E entry vector p3E-IRES-mScarlet-pA. The wild type zebrafish *pik3ca* CDS was introduced into pDonr221 (Invitrogen) using Gateway™ BP Clonase™ II (ThermoFisher Scientific) to make pME-*pik3ca^WT^*. *pik3ca* hotspot mutation sites were introduced into pME-*pik3ca^WT^* sequence by designing back-to-back mutagenesis primers incorporating hotspot mutations *H1048R/L*, *E542K* or *E545K*, phosphorylating them using T4 PNK (NEB) according to manufacturer’s instructions, and amplifying using Q5 polymerase (NEB). T4 ligase (NEB) was used to re-ligate the mutated amplicon, and the original template digested using Dpni (NEB). LR Clonase II+ (ThermoFisher Scientific) was used to combine p5E entry vectors p5E-*β-actin* (54), p5E-*fli1ep* (50), p5E- *tbxta* (this study, based off (55), with a *pik3ca-PROS* pME, and p3E-IRES-mScarletpA (this study, modified from p3E-IRES-GFP) into pDestTol2CG2 (54) which carries a *cmlc2:eGFP* reporter. Sanger sequencing was used to confirm successful mutation of pME-*pik3ca^WT^* to pME- *pik3ca^H1048/H1048L/E542K/E545K^*and correct integration of all components. Sequences of primers used for plasmid construction are found in **Supplementary Table 1**.

### Generation of mosaic *pik3ca^PROS^* fish

Approx. 2 nl of 30 ng/μl transgenesis construct was co-injected with 35 ng/μl Tol2 mRNA into zebrafish embryos at the 1 cell stage. Embryos were *Tg(fli1:eGFP)* to enable visualization of blood vessels, in either a wildtype or *casper* (unpigmented) genetic background – the latter to more easily image embryos beyond stages of melanocyte development. Embryos were screened for successful PROS transgene integration from 30 hpf onwards by assessment of *cmlc2:GFP* cardiac fluorescence and/or *mScarlet* fluorescence. Morphologically abnormal embryos at this stage were discarded, as vascular development is disrupted due to defects in somite and heart development, for example (56). Scoring of vascular malformations was performed at 72 hpf. For studies on older fish, embryos were raised to juvenile stages.

### Imaging

Embryos and juvenile fish were anaesthetized with MS:222 and if necessary, placed on 3% methylcellulose. Fluorescent mesoscope images were acquired using a Leica M165FCA fluorescence stereomicroscope fitted with a 0.63x and 1x Plan-Apochromat objective, and a Leica LMT260 XY scanning stage (Leica Microsystems, Germany). Image acquisition was performed using a Leica K8 monochrome cMOS camera in Leica Application Suite software (LASX; version 3.7). Color images were acquired as above but using a Leica KC3 color cMOS camera (Leica, Microsystems, Germany). Images were acquired in Leica Application Suite software (LASX; version 3.7). Movies were generated using Imaris (v9.6.0). Images were processed using FIJI (v2.9.0).

Confocal images were acquired using a 20X/0.75 lens on the multimodal Imaging Platform Dragonfly (Andor technologies, Belfast UK) equipped with 488 and 561 lasers built on a Nikon Eclipse TI-E inverted microscope body with Perfect focus system (Nikon Instruments, Japan. Data were collected in Spinning Disk 25 μm pinhole mode on the iXon 888 EMCCD camera using a Bin of 1 and no frame averaging, and using Andor Fusion acquisition software. Z stacks were collected using the Nikon TiE focus drive. Movies were generated using Imaris (v9.6.0). Images were processed using FIJI (v2.9.0).

### Bone and Cartilage staining

To stain bone and cartilage in juvenile fish, the Juvenile No Acid Bone & Cartilage Stain was followed as described (zfin.atlassian.net/wiki/spaces/prot/pages/352157857) using a 0.02% Alcian Blue 8GX (Merck)/10 mM MgCl_2_ solution to stain cartilage and then a 0.01% Alizarin red S (Merck) solution to stain mineralized bone.

### FACS and library preparation

*Tg(fli1:eGFP)* embryos were injected with either *tbxta:pik3ca^E545K^-mScarlet* or *tbxta:mScarlet* and >300 embryos per condition raised to 19 hpf. Each batch was dissociated to single cell suspensions using the methods described in (57). Cells from stage matched wild-type embryos were used as negative controls to gate fluorescence. Live cells were sorted using a FACSAria SORP instrument (BD Biosciences UK), with GFP (525/50, BP/488 nm laser), RFP (582/15 BP, 561 nm laser), and DAPI (450/20BP, 405 nm laser) filters into 2% BSA in PBS and immediately processed using the Chromium platform (10x Genomics). Libraries were prepared using v3.1 Standard Throughput kits according to 10x protocols (10x Genomics), and quality control and quantification assays performed using High Sensitivity DNA kits on a Bioanalyzer (Agilent). Libraries were initially sequenced on an Illumina NovaSeq SP Flow cell with 100 cycles with two lanes per sample. The imbalances in cell number/sequencing depth between the two samples necessitated re-sequencing of the PROS sample as before. This resulted in an average of 120,316 reads/cell after normalization.

### Bioinformatics

FastQ files were aligned using the CellRanger (v8.0.1, 10x Genomics) mkref, mkgtf and aggr pipelines to custom zebrafish STAR genome index using annotations from Ensembl GRCz11 release 112 with manual addition of *eGFP* and *mScarlet*. The resultant h5 matrices were read into R (v4.4.2) and subjected to quality control using Seurat (58) guidelines (v5.2.1, features >200, <13000 and mitochondrial gene < 5%). This resulted in 10306 cells from PROS embryos and 3505 from control. Data were scaled and regressed to cell cycle stage before performing PCA, finding the dimensionality of the data by elbow plot (dims=50) to plot 40 clusters on UMAP plots (SeuratExtend (v1.1.4 (59)). SingleR (60) (v2.8.0) was used with previously published datasets GEO: GSE112294 (35) and NCBI SRA PRJNA940501 (36) to call clusters. DE analyses were performed using zinbwave (v 1.28.0 (61)) and DESeq2 (v1.46.0(62)), selecting genes with padj<0.05 before using clusterProfiler (v4.14.3 (63)) to perform gene enrichment analyses searching Reactome, GO:BP and KEGG libraries, filtering by p.adj <0.05 (raw data are in **Supplementary Table 2**). Additional footprint enrichment analyses were conducted using DecoupleR (v. 2.12.0 (64)) using the inbuilt Progeny databases and zebrafish Hallmark datasets imported from MSigdDB according to package guidelines, with Seurat’s FindAllMarkers with wilcox testing to determine changes (p.adj < 0.1, p<0.05) to pathway enrichment. Proportions of cells within cluster for PROS and control data were found by dividing the number of cells per cluster/total cell number per experimental condition, with Pearson’s chi-squared test (two-tailed) used to determine significance (raw data are in **Supplementary Table 3**). RNA velocity analysis was performed on loom files for control and PROS data generated with velocyto (v. 0.17 (65)), and velocity computed using scVelo’s (v.0.2.5 (66)) dynamical modelling approach, and visualized using heatmaps and stream plots.

CellChat (v. 2.1.2 (37)) was used to infer potential signaling events between clusters according to relevant package guidelines. NICHES (v. 1.0.0 (38, 67)) was used to perform ligand-receptor expression analysis at single cell resolution, with differential expression analysis within Seurat, using the MAST test, performed to identify significant changes to LR gene expression (raw data are in **Supplementary Table 4**). Additionally, circuit diagrams were generated using code from https://github.com/RaredonLab/NICHESMethods. Plots were generated using Seurat, CellChat, NICHES, Complex Heatmap (v. 2.22.0 (68)) and ggplot2 (v. 4.0.0).

## Acknowledgments

This work is supported by funding from the CLOVES Syndrome Community [c-13277479] and the UK Medical Research Council [MR/Z506321/1]. We are grateful to Kristen Davies and Seth Haddix from the CLOVES Syndrome Community for discussions about the experimental aims and design of this work, and to Mihaly Badonyi and Joe Marsh (MRC Human Genetics Unit) for assistance with Pymol. RKS is also supported by the BHF Centre for Research Excellence Award III [RE/18/5/34216]. RRM is funded by a UKRI Future Leaders Fellowship [MR/Y017439/1]. MSBR and NW are funded by T32GM086287 from the NIGMS, and laboratory startup funds from the Yale School of Medicine and the Yale Department of Anesthesiology. EEP is funded by the Medical Research Council (MC_UU_00035/13) and the Cancer Research UK Scotland Centre (CTRQQR-2021\100006).

## Data and Code availability

scRNA-seq experiments generated in this study have been deposited at GEO and will become publicly available as of the date of publication. Computational code and plasmid constructs generated for this are available from the lead contact upon request.

## Author Contributions

Conceptualization: HB, RKS, EEP; Data curation: HB; Code checking: RRM, NW, MSBR; Formal analysis: HB; Funding acquisition: RRM, MSBR, RKS, EEP. Investigation: HB; Methodology: HB, RRM, MSBR, EEP; Project administration: RKS, EEP; Supervision: MSBR, RKS, EEP; Visualization: HB; Writing – original draft: HB, RRM, RKS, EEP; Writing – review & editing: All authors

## Supplementary Figure Legends

**Supplementary Figure 1:**
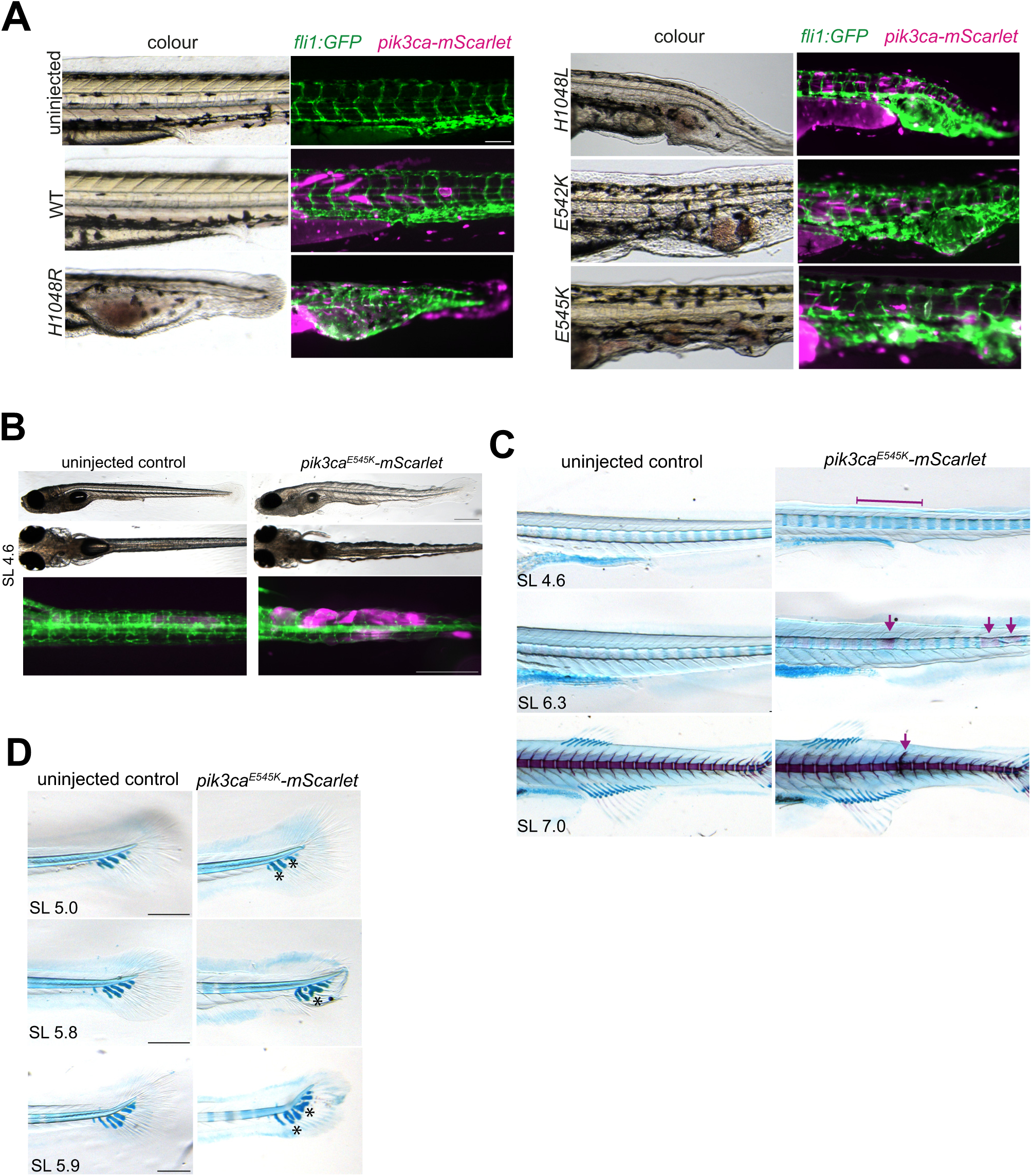
Hotspot *pik3ca^PROS^* transgenes cause vascular malformations and defects in muscle, cartilage and skeletal patterning in larval fish. A) Representative images of severe vascular malformations found in 72 hpf *Tg(fli1:eGFP)* embryos previously injected with either *β-actin:pik3caH1048R*, *H1048L, E542K* or *E545K.* Uninjected embryos, and those injected with WT *pik3ca* showed no phenotype at this timepoint. In *pik3ca^PROS^* embryos, colour panels on the left show blood pooling in malformed vessels, with matched merged fluorescence images of the same area on the right showing enlarged *fli1:eGFP+* vessels (green), and variable numbers of mosaic *pik3ca^PROS^-mScarlet* cells (magenta). Scale bar = 100 μm. B)*Tg(fli1:eGFP);casper* fish injected with *β-actin:pik3ca^E545K^-mScarlet* but negative for vascular malformations at 72 hpf developed a ‘rippled’ appearance by two weeks of age (SL4.6) caused by protrusions of *pik3ca^PROS^-mScarlet* cells in the embryo flank (10/25). Scale bar = 100 μm. C) Patterning defects in the developing spine of staged matched *Tg(β-actin:pik3ca^E545K^-mScarlet; fli1:eGFP); casper* injected embryos, showing irregular spacing of cartilage centra during early spine development (3/9 fish in which pattern had developed, prior to bone deposition), premature deposition of bone in patches posterior to stage-matched controls (1/3 fish at a stage where bone deposition is just starting in anterior and so therefore premature deposition is noticeable), and thickened vertebra structure (1/6 fish in which vertebrae were formed) D) Comparison of stage-matched uninjected and *Tg(β-actin:pik3ca^E545K^-mScarlet; fli1:eGFP); casper* injected fish during tail fin development with patterning defects, revealed by alcian blue staining for cartilage and alizarin red co-staining for bone. 3/9 cartilage hypural patterning defects, 5/7 tail shape defects.

**Supplementary Figure 2:**
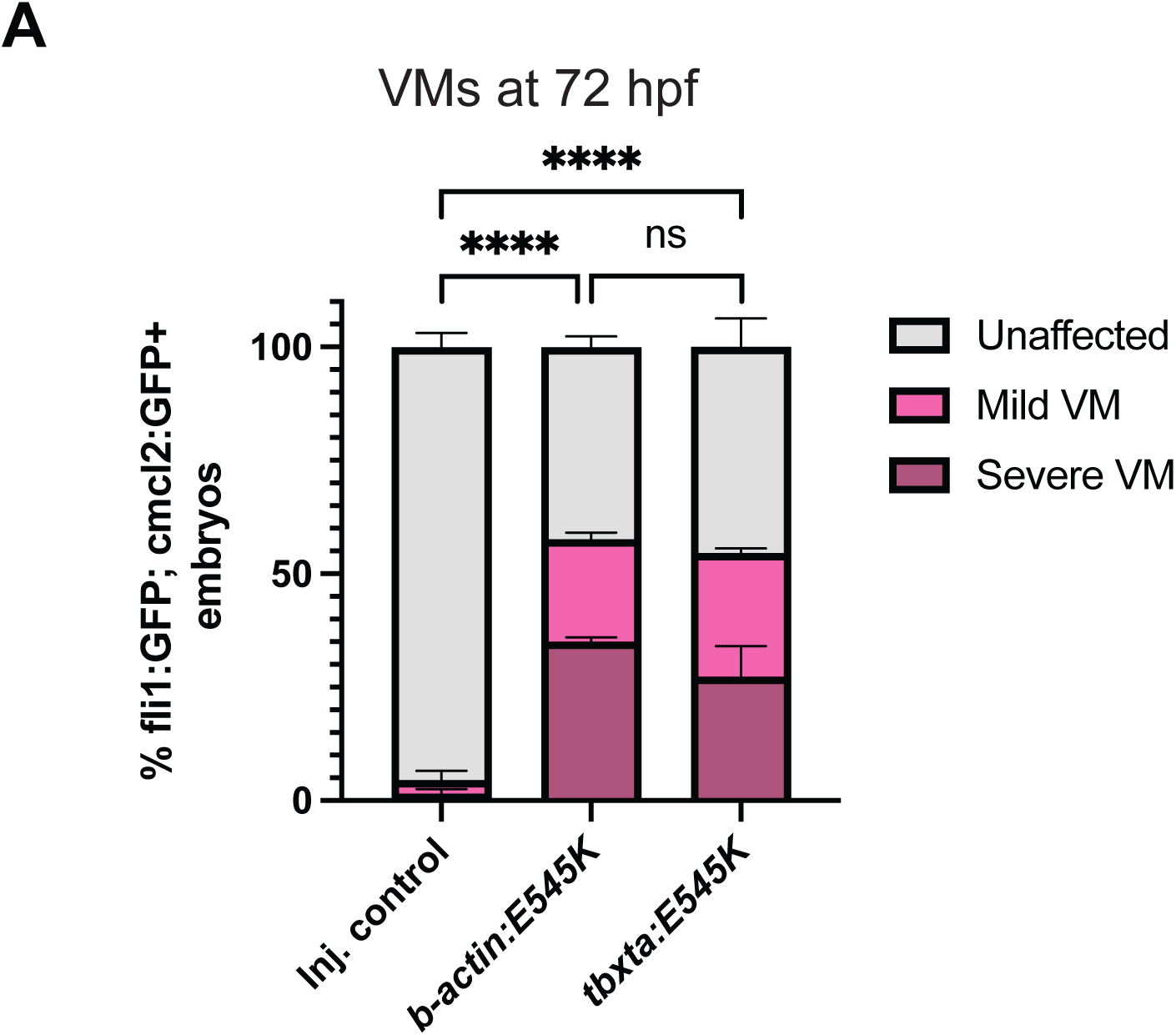
Ubiquitous and mesodermal expression of *pik3ca^PROS^* causes vascular malformations. A) Quantification of vascular phenotypes in 72 hpf Tg(*fli1:eGFP*) embryos injected with *pik3ca^E545K^-mScarlet* constructs driven by *β-actin* or *tbxta*, or an injected no-*pik3ca* control (*tbxta:mScarlet*). Represents 3 biological replicates. Error bars are S.E.M. p-values are from chi:squared tests on raw counts, * p<0.05, ***p<0.001, **** p<0.0001.

**Supplementary Figure 3:**
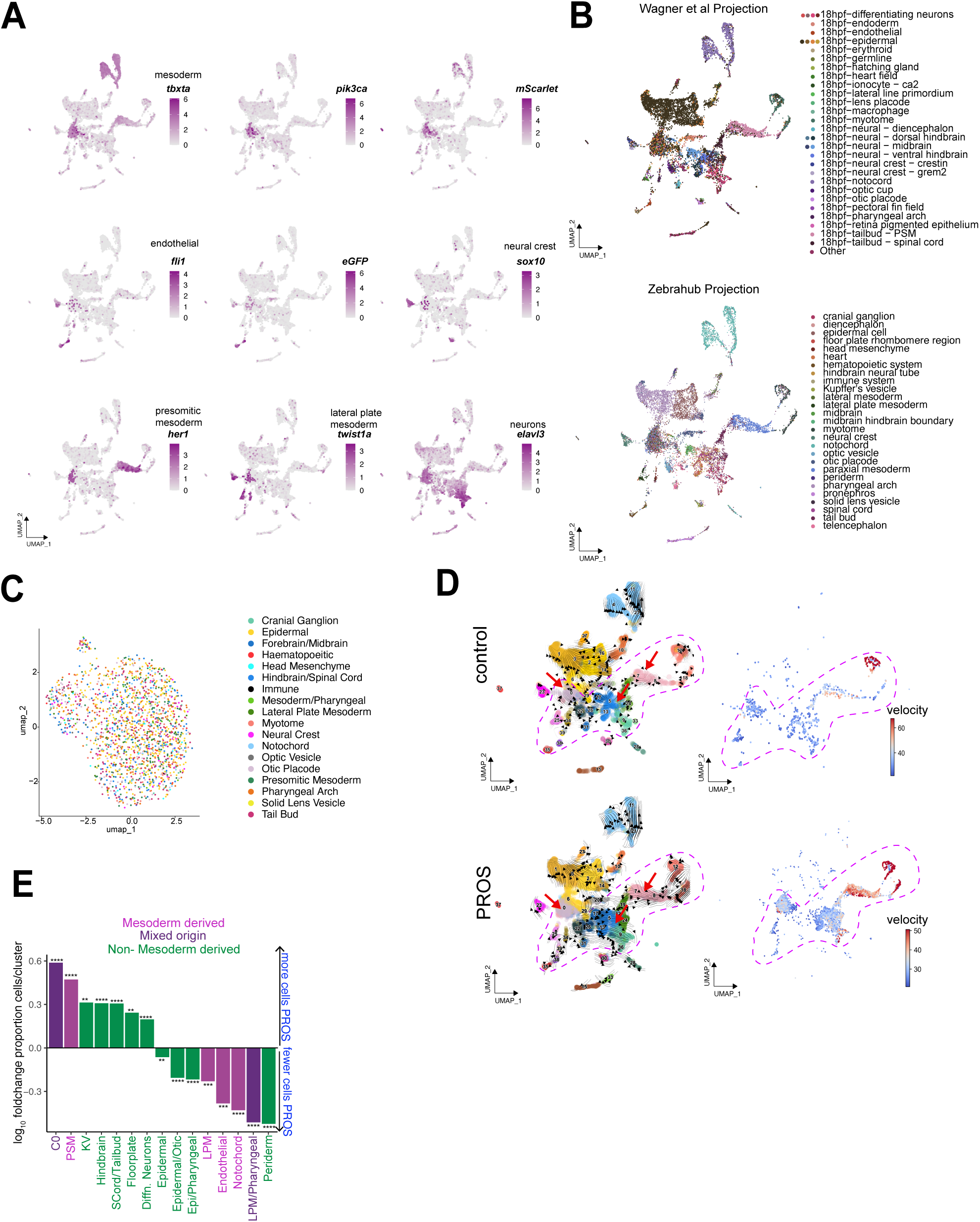
scRNA seq analysis of *pik3ca^PROS^* mosaic embryos. A) UMAP FeaturePlots showing expression intensity (purple) of transgenesis or cell lineage markers. B) Mapping of 18hpf (top) and 19 hpf (bottom) published dataset identities onto these data to call clusters. C) UMAP of re-clustered cluster 0, with identities from the zebrahub 19 hpf dataset mapped as in B. D) Single cell velocity analysis was performed on PROS and control data, and the vector field overlaid on a UMAP projection, with arrow indicating velocity dynamics and strength across clusters. The greatest variance in arrow length and density between PROS and control is found in the cluster groups outlined (pink dashed). The velocity length of this subset is displayed on a heatmap, with cells showing the highest abundances of unspliced mRNA in red. E) As Figure 4E, but to show the extent of the statistically significant changes in the proportion of cells within cluster groups. Two-sided Pearson’s chi-squared proportion test. ** p<0.01, ***p<0.001, ****p<0.0001

**Supplementary Figure 4:**
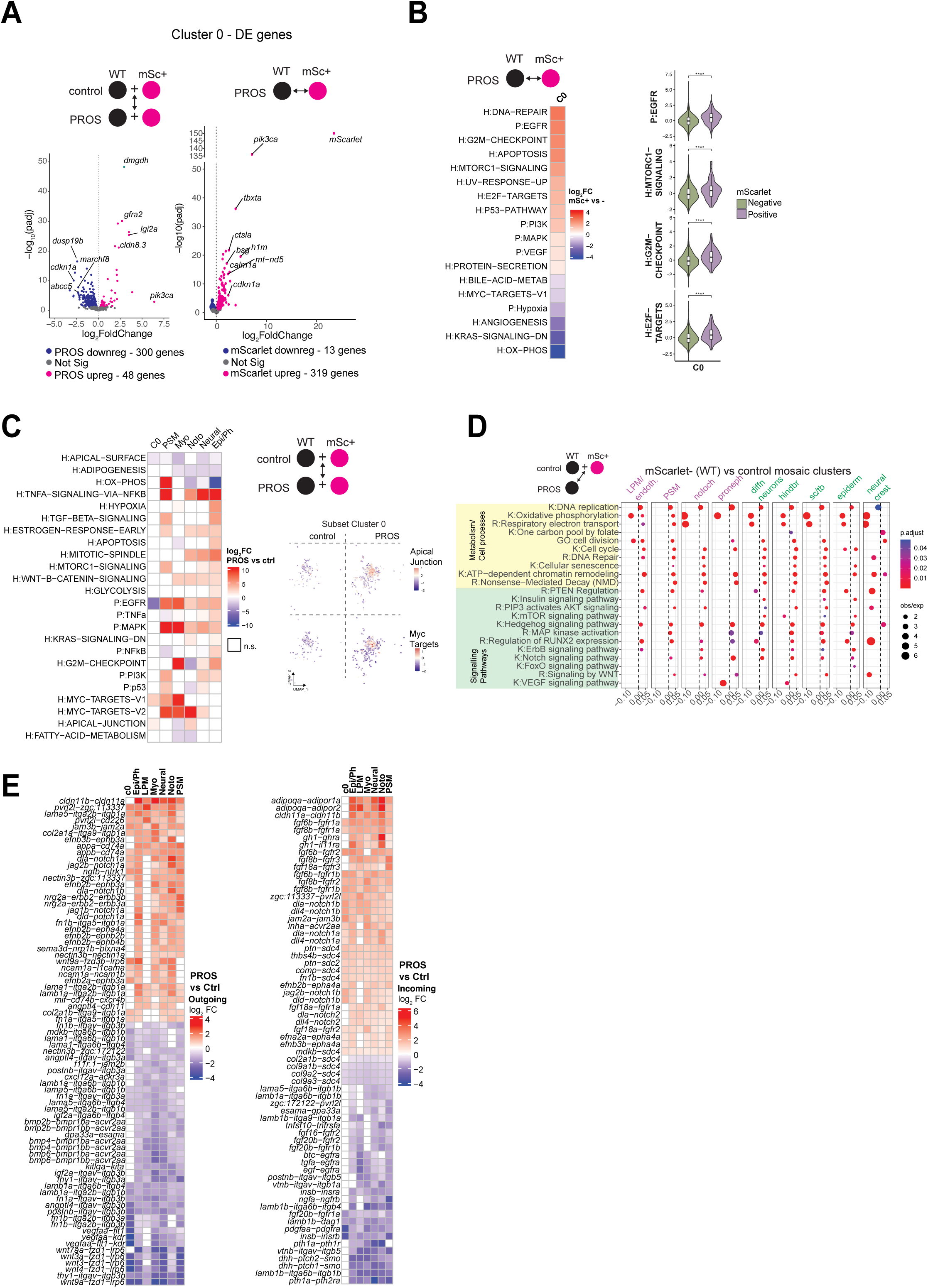
Mosaic *pik3ca^PROS^* causes pan-lineage transcriptional changes, including to ligand-receptor signalling genes. A) Volcano plots showing significantly differentially expressed genes after DE analysis of cells within cluster 0. Left – comparison of all cluster 0 PROS mosaics vs control, Right – mScarlet+ vs mScarlet-cells in PROS mosaics only. B) Heatmap showing fold change enrichment (Seurat wilcox test p<0.05, p.adj<0.1) of Hallmark (H:) and Progeny (P:) pathways in PROS mosaic cluster 0 mSc+ vs mSc-cells. Violin plots confirm significant upregulation of selected Hallmark signatures in mScarlet+ cells (wilcox test, **** = p<0.001) C) Heatmap showing fold change enrichment (Seurat wilcox test p<0.05, p.adj<0.1) of Hallmark (H:) and Progeny (P:) pathways in PROS mosaics relative to control N.S = non-significant, and Cluster 0 UMAP subset showing expression intensity of Hallmark terms, split by experimental condition. D) As in Figure 4F), but comparing only *mScarlet-* (ie, WT) cells between experimental conditions. E) Heatmaps depicting log_2_ fold change in expression of the top differentially expressed “sending” (left) and “receiving” (right) ligand-receptor gene pairs in PROS relative to control embryos, clustered by row. Only differences for which p.adj<0.05 by Wilcoxon test between PROS and WT are shown.

## Movie Legends

Movie 1-3:

Movie 1 – Maximum projection movie of Figure 4A of uninjected *Tg(fli1:eGFP)* embryo showing the arrangement of vessels around the central notochord.

Movie 2 - Maximum projection movie of a Mild vascular lesion in a *Tg(fli1:eGFP)* embryo injected with *pik3ca^E545K^-mScarlet* (see Figure 4B) showing first separated *pik3ca^PROS^ mScarlet* expression (magenta) and then merged with *fli1:eGFP*

Movie 3 – Maximum projection movie of a Moderate vascular lesion in a *Tg(fli1:eGFP)* embryo injected with *pik3ca^E545K^-mScarlet* (see Figure 4B) showing first separated *pik3ca^PROS^ mScarlet* expression (magenta) and then merged with *fli1:eGFP.* Cells co-expressing *GFP* and *mScarlet* are white

Movie 4 – Timelapse of injected mosaic mosaic *tbxta:mScarlet Tg(fli1:eGFP)* control embryo between 16-28.5 hpf, showing normal development of blood vessels, with ISV sprouting and DLAV formation from anterior (left) to posterior (right).

Movie 5 – Timelapse of injected mosaic *pik3ca^PROS^ Tg(fli1:eGFP)* embryo between 16-28.5 hpf showing ectopic *fli1:eGFP+* endothelial cells contributing to premature formation of the DLAV, and defects in ISV sprouting.

## Supplementary Tables

Supplementary Table 1 – Primers used for plasmid construction

Supplementary Table 2 – Raw results from Deseq2 DE analyses used to generate Figure 4F

Supplementary Table 3 – Raw values used to generate proportion plots in Figures 4E and Supplementary Figure 3E.

Supplementary Table 4 – Raw results from DE analyses on NICHES signalling objects used to generate Figure 5C

